# GDF15 regulates necroptotic cell death through direct interaction with RIPK3

**DOI:** 10.1101/2025.10.23.684070

**Authors:** Anastasia Piskopou, Isabella Vinson, Kong Sit, Els Sweep, Maarten Altelaar, Kelly E. Stecker

## Abstract

Enhancing the immunogenicity of tumor cells is a major objective in cancer therapy, particularly for tumors with low immune cell infiltration scores. Inducing immunogenic forms of programmed cell death (PCD) offers a promising strategy to strengthen anti-tumor immunity and improve therapeutic outcomes. Necroptosis, a highly inflammatory form of regulated cell death triggered by TNF signaling, can elicit robust immune activation. However, its regulation in tumor cells remains incompletely understood, limiting its therapeutic exploitation. To investigate the protein-protein interactions that govern necroptotic cell death in tumor cells, we established a co-Immunoprecipitation - Mass Spectrometry (coIP-MS) workflow using RIPK3, the central effector kinase driving necroptosis in TNF-induced signaling, as bait. This unbiased proteomic approach enables the identification of candidate regulators directly associated with the necrosome complex components under active necroptotic conditions. Among identified candidates, TCOF1 and GDF15 emerged as previously unrecognized modulators, with functional knockout of either gene markedly enhancing necroptotic cell death in tumor cells. Reciprocal IP experiments confirmed a direct interaction between GDF15 and RIPK3, supporting its mechanistic role as a negative regulator that suppresses necroptotic signaling. Thus, our findings extend the function of GDF15 beyond its established role in inflammation, uncovering an additional layer of regulation at the level of cell-intrinsic death signaling. Collectively, our findings position GDF15 as a RIPK3-interacting “brake” on necroptotic cell death and highlight TCOF1 as an additional inhibitory node. Our study underscores the potential of targeting necroptosis-suppressive mechanisms to influence PCD outcomes in tumors and demonstrates the power of coIP-MS for mapping TNF-induced interactions to reveal actionable molecular targets for tumor sensitization.

## INTRODUCTION

In recent years, therapies that target inhibitory immune pathways have reshaped the clinical landscape of oncology. Immune checkpoint blockade (ICB) represents the most prominent of these approaches, acting through blockade of receptors such as PD-1 or CTLA-4 to restore cytotoxic T-cell activity and achieve durable tumor control in a subset of patients^1–3^. However, ICB is not uniform across cancers and depends strongly on the tumor microenvironment (TME)^4^. Tumors are broadly categorized into “hot” (immune-inflamed), “variable” (immune-excluded), and “cold” (immune-desert) phenotypes based on immune cell infiltration^4,5^. Hot tumors, enriched in cytotoxic CD8⁺ T cells within the tumor parenchyma, respond favorably to ICB. In contrast, cold tumors, which lack meaningful immune infiltration, are frequently resistant^6,7^. This highlights the need for strategies that can convert non-inflamed tumors into immunologically active ones by increasing their susceptibility to immune attack.

One promising approach is to induce immunogenic forms of programmed cell death (PCD), which can release tumor antigens and inflammatory mediators that recruit and activate immune cells^8,9^. Among these, necroptosis has attracted significant interest as it is a lytic, pro-inflammatory cell death pathway that potently stimulates innate and adaptive immune responses^10,11^. Unlike apoptosis, which is non-inflammatory and involves orderly cellular breakdown without membrane rupture, necroptosis results in cell swelling, plasma membrane disruption, and the release of damage-associated molecular patterns (DAMPs). DAMPs are then sensed by pattern recognition receptors (PRRs) on dendritic cells and macrophages, initiating robust inflammatory cascades and T-cell priming^12,13^.

Necroptosis can be triggered by several upstream signals, including Toll-like receptors, Z-DNA binding protein 1 (ZBP1), and interferon receptors, but the TNF–TNFR1 axis represents its prototypical activation pathway^14–17^. TNFR1 engagement by TNFα initiates the assembly of a membrane-bound complex (complex I) containing TNFR1-associated death domain protein (TRADD), receptor-interacting protein kinase 1 (RIPK1), TNFR-associated factor 2 (TRAF2), linear ubiquitin chain assembly complex (LUBAC), and cellular inhibitors of apoptosis (cIAP1/2). Canonical downstream TNFR1 signaling, typically promotes cell survival through activation of NF-κB and MAPK pathways, however this outcome is highly dictated by the activation state of RIPK1 which acts as a molecular switch^18,19^. Upon activation, the survival signal is bypassed, and RIPK1 transitions into a cytosolic death-signaling platform promoting the assembly of apoptosis- or necroptosis-inducing complexes. This switch is tightly regulated by post-translational modifications such as ubiquitination and phosphorylation, which act as checkpoints balancing survival and cell death^20–22^. Once these checkpoints are overcome, RIPK1 engages RIPK3 through RIP homotypic interaction motifs (RHIM), forming an amyloid-like complex known as the necrosome complex^23^. RIPK3 is a key effector kinase that orchestrates necroptotic signaling, functioning as the central hub of the pathway, receiving input from RIPK1 and amplifying the death signal through its own phosphorylation^23,24^. Once the necrosome complex is formed, RIPK1 and RIPK3 both undergo activating phosphorylation, resulting in the phosphorylation of mixed-lineage kinase domain-like protein (MLKL). Activated MLKL oligomerizes, translocate to the plasma membrane, and forms disruptive pores, leading to cell lysis and DAMP release^11,25^. MLKL thus serves as the terminal effector of necroptosis and a critical determinant of its immunogenic potential.

While the molecular framework of TNF-induced necroptosis has been extensively studied, its regulation in tumor cells and particularly the identity of modulators that restrain or fine-tune necrosome assembly and cell death execution, remains incompletely understood^23,26,27^. Such regulators could represent potential therapeutic targets for sensitizing resistant tumors to immune attack by lowering their threshold for necroptotic cell death. To address this gap, we employed an unbiased co-Immunoprecipitation–mass spectrometry (coIP-MS) strategy using RIPK3 as bait to map protein-protein interactions under active necroptotic signaling in tumor cells.

## RESULTS

### TNF/Smac/zVAD treatment of HT-29 cells serves as a robust model to study necroptosis

Necroptosis is tightly controlled by upstream checkpoints and frequently suppressed in cancer cells, making it challenging to study under physiological conditions^28,29^. To explore the molecular machinery of the necrosome and necroptotic signaling, it is essential to establish a robust model of necroptosis induction in tumor cells. To achieve this aim, we selected HT-29 colon carcinoma cells, which are known to be sensitive to classic necroptosis inducers^30,31^.Cells were treated with a combination of TNFα, the Smac mimetic Birinapant, and the pan-caspase inhibitor zVAD-FMK (hereafter TBZ treatment), a combination widely used to trigger necroptotic cell death^31^. Biochemical validation demonstrated robust phosphorylation of RIPK1 (Ser166), RIPK3 (Ser227), and MLKL (Ser358), consistent with activation of the canonical necrosome cascade under necroptotic stimulation (Fig. 1B). We next used the Incucyte SX1 live-cell imaging system to monitor treated HT-29 cells over 24 hours. As expected, treatment with TNFα alone did not induce detectable cell death in HT-29 cells (Fig. 1C and D), indicating that additional modulators are required to overcome intrinsic survival mechanisms. In contrast, the combination treatment TBZ, efficiently triggered cell death, which was fully rescued by the RIPK1 kinase inhibitor necrostatin-1 (Fig. 1C and D), confirming that the observed phenotype was necroptotic rather than apoptotic.

**Figure 1.**
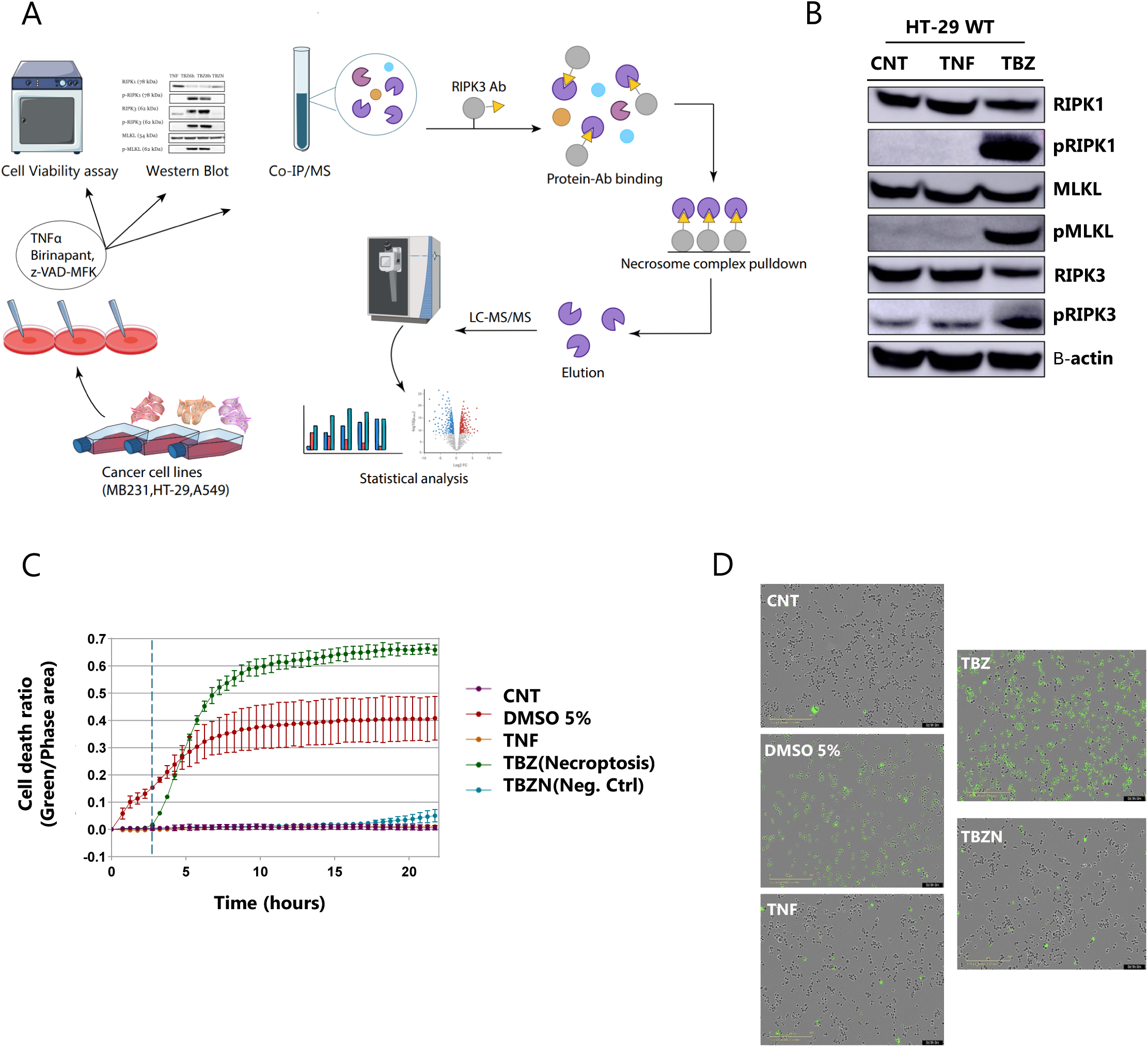
Combination treatment induces necroptotic cell death in HT-29 cells. **(A)** Overview of the experimental workflow. **(B)** Biochemical validation of necrosome formation upon combination treatment in HT-29 cells. Tumor cells were harvested and blotted against the known necroptosis executors. Phosphorylation of RIPK1, RIPK3, and MLKL confirm necrosome complex assembly. B-actin was used as a loading control. **(C)** Live-cell imaging of necroptotic death kinetics under indicated treatments using Incucyte SX1 (Sartorius). Combination treatment TBZN served as a negative control for necroptotic cell death, as Nec-1 inhibits RIPK1 kinase activity, thereby preventing necrosome complex assembly. Tumor cells were scanned in 15-minute intervals over 24 hours post treatment. Error bars represent standard deviation (SD) between replicates (n=3). Background signal was subtracted at the time of treatment (t=0) and green (cell death) signal is represented as a ratio to phase signal. Blue dashed line indicates the onset of necroptotic cell death and timepoint for co-IP LC-MS analysis. **(D)** Representative images of HT-29 cells following 9 hours of treatment. Scale bars: 400 µm.

To test whether this mechanism was conserved across tumor types, we extended our analysis to the triple-negative breast cancer cell line MDA-MB231 (Fig S1A, B). Similar biochemical and functional outcomes were observed, with HT-29 cells displaying higher overall sensitivity to necroptotic death (Fig. S1C). Together, these results establish a tractable necroptotic model in human tumor cells in which we can study the molecular patterns and protein-protein interactions that govern necroptotic signaling.

### coIP–MS workflow maps RIPK3-associated proteins under necroptotic conditions

To identify novel regulators of necroptosis, we established a co-Immunoprecipitation– Mass Spectrometry (coIP–MS) workflow using RIPK3 as bait. HT-29 colon carcinoma cells were treated under the same conditions used for our phenotypic assays and for the TBZ treatment, and incubation was stopped just before necroptotic cell death was initiated (Fig1C). RIPK3 immunoprecipitations were performed in parallel with a nonspecific IgG antibody as a negative control. Principal component analysis confirmed a clear separation of TBZ-treated RIPK3 pulldowns from the controls, indicating distinct protein complex formation under necroptotic conditions (Fig. S2A). RIPK3 pulldowns under TBZ treatment reliably captured known necrosome interactors, including RIPK1, CASP8, and FADD (Fig. 2A). The enrichment of these canonical partners was expected, as TBZ treatment stabilizes the necrosome by preventing caspase-mediated cleavage and allowing RIPK1–RIPK3 assembly^32^. Notably, pathway analysis of enriched proteins in TBZ-treated IP samples showed strong enrichment for the necroptotic signature, alongside proteasome-associated signatures. This may reflect protein turnover due to ongoing necroptotic signaling and antigen processing linked to necroptotic death (Fig. 2B). When comparing RIPK3 complexes in TBZ vs TNF treatment alone, we observed an even more specific enrichment of necroptotic signatures (Fig. 2C). Importantly, RIPK1 recruitment to RIPK3 was markedly reduced in the TNF-only condition, underscoring that full necrosome assembly requires caspase inhibition and IAP antagonism. By contrast, TBZ treatment strongly promoted RIPK1-RIPK3 interactions, consistent with our phenotypic and biochemical validation of necroptosis induction (Fig. S2D).

**Figure 2.**
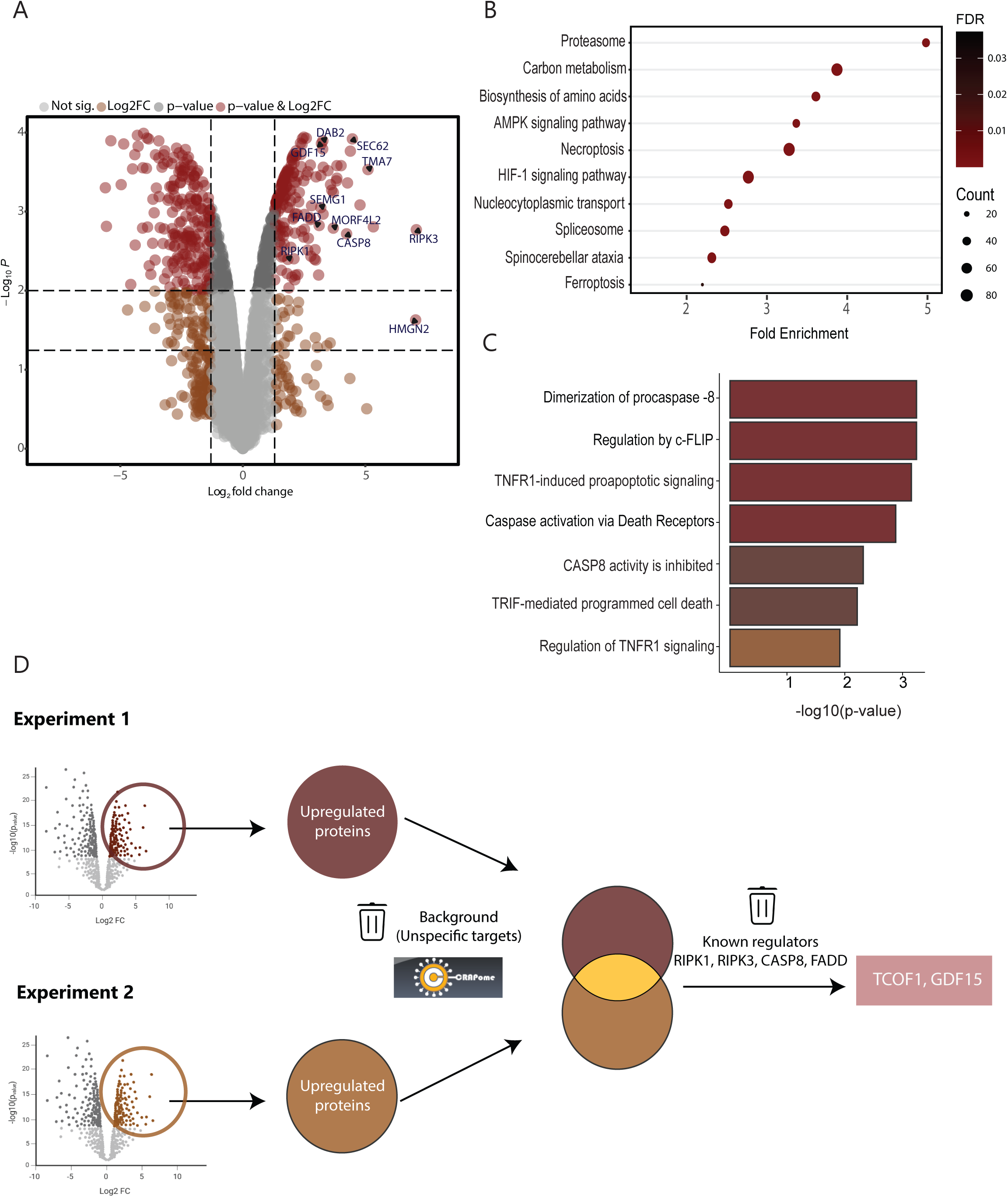
coIP-MS workflow maps RIPK3-associated proteins under necroptotic conditions. **(A)** Relative abundance of TBZ-induced RIPK3 pull downs against IgG Ctrl. Dashed lines indicate cutoffs at p < 0.05 and p < 0.01. **(B)** KEGG pathway enrichment of significantly enriched interactors upon combination treatment, showing strong enrichment for necroptosis, proteasome, and metabolic pathways (FDR < 0.05). **(C)** Reactome analysis of enriched proteins relative to TNF treated cells, highlighting cell death–related pathways, including TNFR1-induced proapoptotic signaling and caspase-8 inhibition. **(D)** Strategy for filtering high-confidence interactors across replicates. Two independent experiments (total n = 4) were compared, retaining proteins significantly enriched or uniquely present under TBZ treatment. Background proteins and contaminants were removed using the CRAPome (2.0) database for affinity purification. Interactors also detected in the non-specific IgG controls were excluded.

To identify novel interactions that govern necroptotic signaling, we repeated the pulldown experiments in additional independent biological replicates, resulting in a total of four necrosome co-immunoprecipitations performed across two experimental batches. Upregulated proteins were compared across experiments, and common interactors were retained as candidate hits, while proteins found in the IgG control or classified as contaminants by using the CRAPome database were filtered out (Fig. 2D)^33^. This stringent analysis reduced background noise and allowed us to focus on high-confidence RIPK3-associated proteins.

Through this workflow, we identified both established regulators of necroptotic signaling, including RIPK3, RIPK1, CASP8, FADD, and proteins not previously linked to cell death. Among the latter, we identified two candidates that reproducibly emerged across independent experiments: TCOF1 and GDF15 (Fig 2A, Fig S2C). While GDF15 is recently recognized as an immune-modulatory cytokine (^34,35^), neither protein has been described in the context of necroptosis, suggesting they may represent previously unrecognized modulators of RIPK3-dependent cell death signaling.

### TCOF1 and GDF15 knockout enhances necroptotic cell death in HT-29 cells

To test the functional relevance of candidate interactors identified by our coIP–MS workflow, we focused on TCOF1 and GDF15, which emerged as novel putative regulators of necroptosis. TCOF1 encodes the nucleolar phosphoprotein Treacle, a key factor in ribosomal DNA transcription and ribosome biogenesis^36,37^. Beyond its canonical role in nucleolar homeostasis, TCOF1 has been implicated in the cellular stress response, where it coordinates DNA damage repair and interacts with p53 signaling pathways^38,39^. Loss of TCOF1 can sensitize cells to oxidative and genotoxic stress, suggesting a broader role in stress tolerance^38^. Given that necroptosis is strongly modulated by stress checkpoints and ribosome-derived signals, we hypothesized that TCOF1 could influence the threshold for necroptotic activation.

GDF15 (growth differentiation factor 15) is a stress-induced cytokine belonging to the TGF-β superfamily^40^. It is transcriptionally upregulated in response to mitochondrial stress, DNA damage, hypoxia, and inflammatory cues^40,41^. Functionally, GDF15 acts as a context-dependent modulator of cell fate and immune signaling: in some settings it promotes tumor growth and immune evasion, while in others it has been linked to apoptosis or metabolic dysfunction^40–42^. Elevated GDF15 levels are frequently observed in cancer and are closely associated with the development of cachexia; however, its specific role in regulated cell death pathways has not been explored in depth^43,44^. Its ability to shape stress and inflammatory responses positioned it as a compelling candidate for regulating necroptotic signaling.

To investigate the role of these proteins in PCD, we generated CRISPR-Cas9 knockouts for both candidates in HT-29 cells (Fig. 3A, 3B). Following TBZ treatment, both TCOF1 KO and GDF15 KO cells displayed significantly accelerated and amplified cell death compared to SgRNA controls (Fig 3C). Time-lapse imaging showed earlier onset of necroptotic cell death, and quantitative analysis of green (cell death)/phase ratios confirmed a significantly higher proportion of dead cells at early time points (1–4 h) in KO cells (1.5–2-fold higher than control, *p* < 0.05). This difference persisted through later stages (5–8 h), where both KO lines maintained consistently higher death rates than sgRNA controls (Fig. 3C), indicating that loss of TCOF1 or GDF15 sensitized tumor cells to necroptotic cell death.

**Figure 3.**
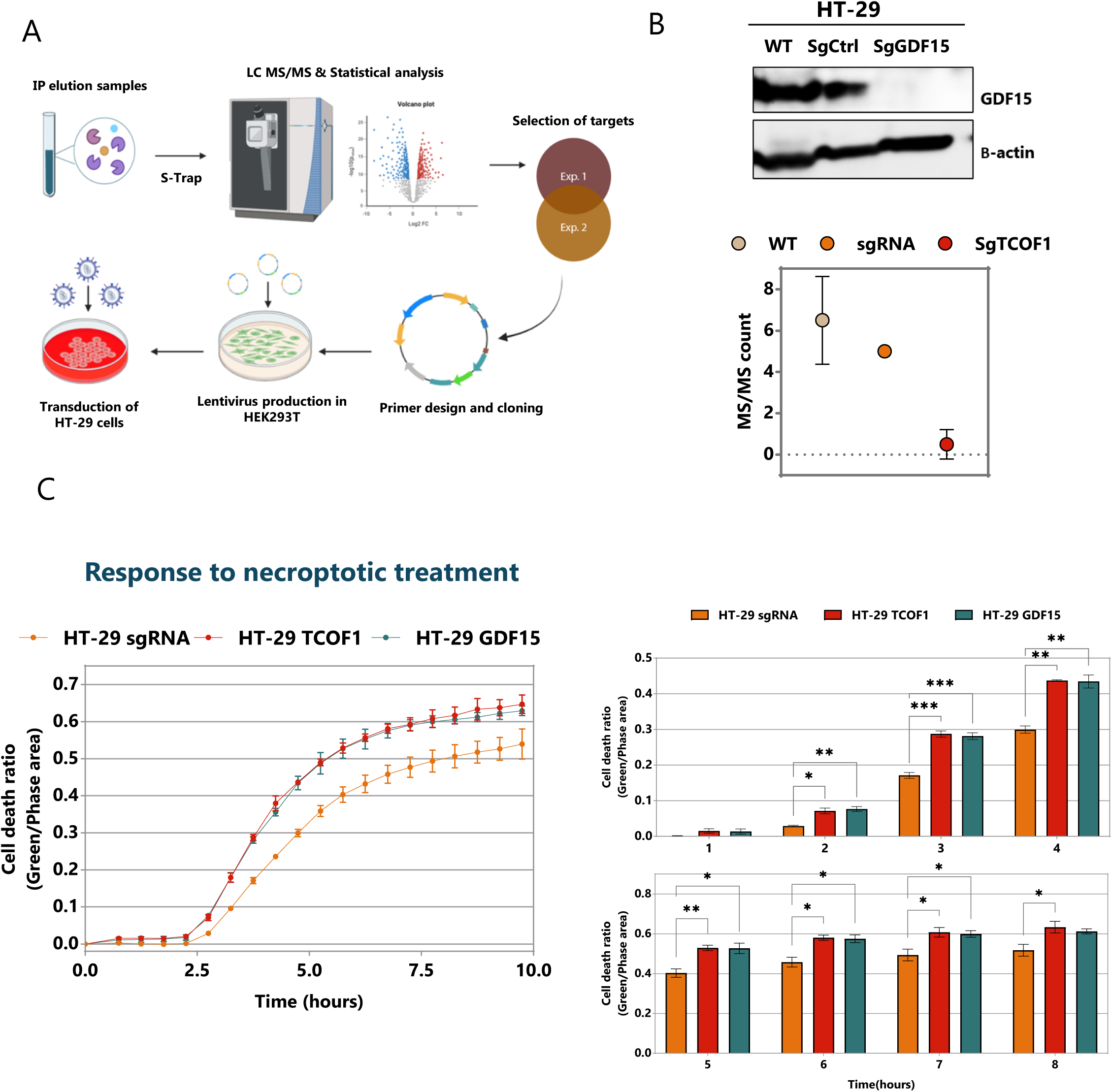
TCOF1 and GDF15 knockout enhances necroptotic cell death in HT-29 cells. **(A)** Overview of experimental workflow for candidate validation. **(B)** Validation of functional GDF15 knockout by Western blot (top) and quantification of MS/MS counts for WT, sgRNA control, and sgTCOF1 cells (bottom). **(C)** Functional validation of TCOF1 and GDF15 KO induced phenotypes by live-cell imaging under TBZ treatment. Left: representative cell death kinetics (green/phase area ratio) measured over 10 h. Right: bar plots showing significantly higher cell death in KO lines compared to SgRNA Ctrl across early (1–4 h) and late (5–8 h) phases necroptotic signaling phases. Data are shown as mean ± SD. Statistical significance was assessed by two-way ANOVA; *p < 0.05, **p < 0.01, ***p < 0.001. Background signal was subtracted at the time of treatment (t=0) and green (cell death) signal is represented as a ratio to phase signal.

Our findings position TCOF1 and GDF15 as novel inhibitory nodes within the necroptotic pathway in HT-29 cells. The underlying mechanisms may differ: TCOF1, through its role in nucleolar integrity and DNA damage signaling, may normally buffer stress responses that converge on necroptotic activation. GDF15, as a cytokine involved in inflammation and tissue homeostasis, may dampen pro-inflammatory signaling cascades that amplify necroptosis, reflecting its complex role in immune regulation. In both cases, their loss appears to removes a layer of restraint, resulting in enhanced RIPK3-dependent signaling and accelerated necroptotic death.

### GDF15 directly interacts with RIPK3 to suppress necroptotic signaling

Given its known association to inflammation, we focused our subsequent investigation on GDF15^34,40^. In recent years, GDF15 has attracted increasing attention as a pleiotropic cytokine with roles beyond its canonical function in metabolism and tissue homeostasis. A growing body of research links GDF15 to inflammation, immune regulation, and stress adaptation across diverse pathological contexts, including cancer, cardiovascular disease, and infection^39,40,42^. This expanding connection to inflammatory processes makes GDF15 a particularly compelling candidate for further study in the context of necroptosis, a form of cell death inherently coupled to inflammatory signaling.

To validate the interaction between GDF15 and RIPK3, we performed reciprocal immunoprecipitation experiments using GDF15 as bait (Fig. 4A), and a non-specific IgG isotype as negative control. In HT-29 cells, GDF15 pulldowns consistently recovered RIPK3, confirming a direct interaction between the two proteins (Fig. 4B). Analysis of whole-cell lysates further revealed a significant upregulation of GDF15 under TBZ treatment, hinting at its involvement in regulated cell death (Fig. 4B). To investigate whether this mechanism was conserved in other cell types, we examined MDA-MB231 cells. In this context, inhibition of GDF15 or TCOF1 also increased necroptotic cell death, although the effects were less pronounced than in HT-29 cells (Fig. S3B). After 24 hours, MDA-MB231 KO cells showed higher overall death rates, but the onset of necroptosis was not accelerated, consistent with its lower baseline sensitivity compared to HT-29 (Fig. S1C). This difference likely reflects the presence of multiple, layered molecular mechanisms that repress necroptosis in MDA-MB231 cells, thereby dampening the impact of single gene perturbations. Importantly, GDF15 pulldowns in MDA-MB231 cells confirmed both GDF15 upregulation under TBZ treatment and its direct interaction with RIPK3 (Figure 4C). Despite the subtler phenotype in this model, these findings further support the hypothesis that GDF15 regulates RIPK3-dependent necroptotic signaling through direct binding to RIPK3.

**Figure 4.**
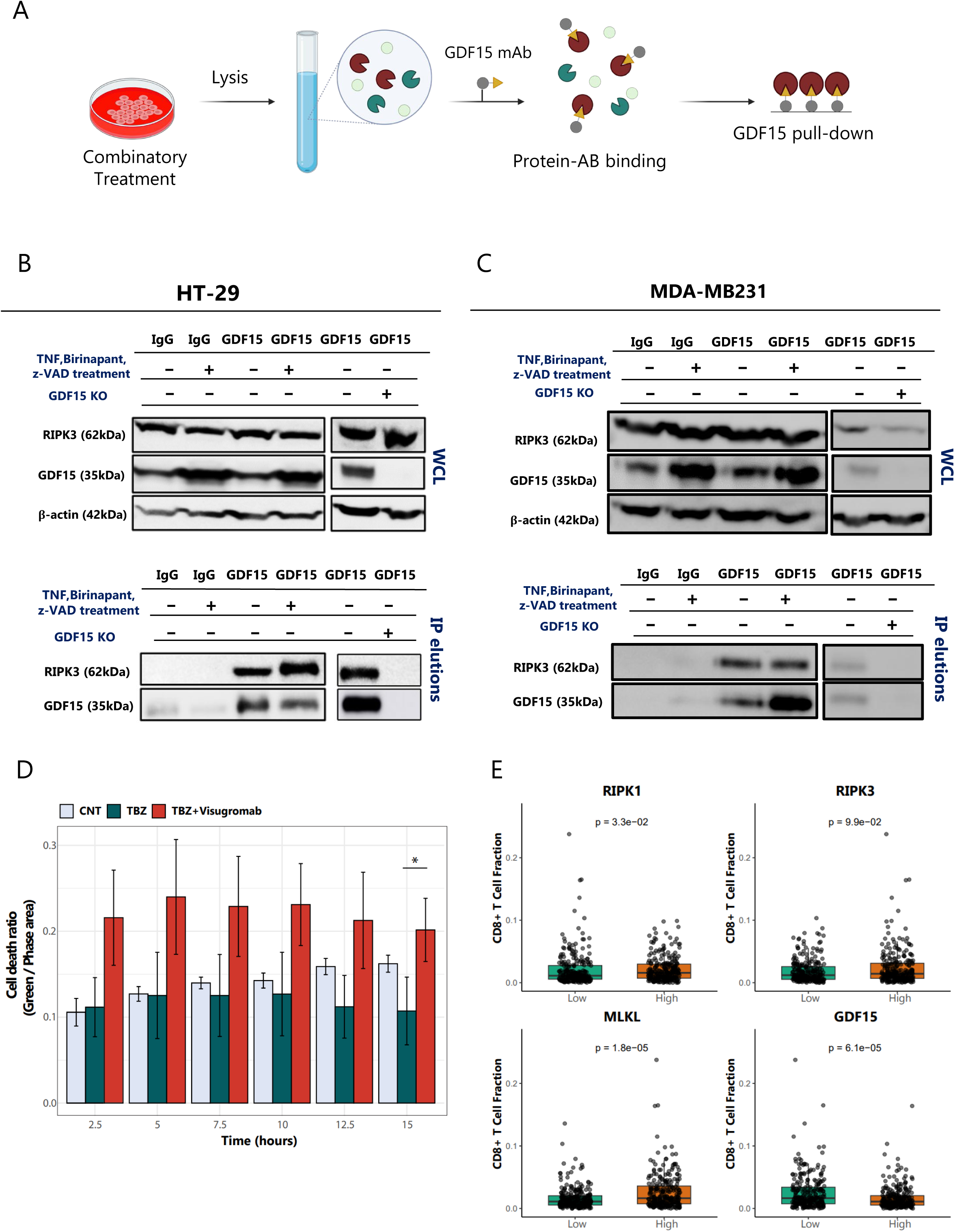
GDF15 interacts with RIPK3 and modulates necroptotic signaling. **(A)** Schematic of GDF15 immunoprecipitation (IP) workflow. **(B)** Reciprocal IP validation in HT-29 cells. Immuno-blotting of whole-cell lysates (WCL) show GDF15 upregulation upon TBZ treatment. B-actin served as loading control. GDF15 pulldowns (IP elutions) recovered RIPK3, confirming direct interaction. **(C)** Same validation experiment performed in MDA-MB231 cells. **(D)** Visugromab neutralization of GDF15 enhances necroptotic signaling in A549 cells. Cell death was quantified as green/phase area ratio using Incucyte over 15 h (mean ± SD, n=3). Statistical significance was assessed by unpaired two-tailed *t*-test; *p*<0.05. Background signal was subtracted at the time of treatment (t=0). Titration of Visugromab in A549 cells showed no significant effects on cell viability in the absence of other treatments (Supplementary Figure 3D). **(E)** Meta-analysis of recently published study employing Lung Adenocarcinoma TCGA tumors, showing correlation between expression of necroptosis regulators and CD8^+^T cell fractions^34^. High RIPK1, RIPK3, and MLKL expression correlated with increased T cell infiltration, whereas high GDF15 expression correlated with reduced infiltration. Values were divided into high and low expression groups by median split and T cell infiltration was compared between groups. Error bars indicate standard deviation between individual measurements (n=142).

We next extended our analysis to A549 lung carcinoma cells, a tumor model previously shown to be resistant to classic necroptosis inducers^31^ (Fig S3C). To explore the effects of GDF15 in this cell line we employed Visugromab, a recently developed neutralizing antibody against GDF15 that has shown to increase treatment efficacy when combined with immune checkpoint blockade in solid tumors^34^. Titration experiments of Visugromab alone on both HT29 and A549 showed no significant effect on cell viability compared to the untreated controls (Fig S3D). Interestingly, however, TBZ treatment in combination with Visugromab significantly increased cell death in A549 cells relative to TBZ alone (Fig. 4D, S3E). These findings suggest a model in which necroptotic signaling induces cellular stress and triggers GDF15 secretion, which then feeds back in an autocrine manner to dampen inflammation and restrain necroptosis. By blocking this feedback loop, Visugromab may remove a critical brake on necroptotic signaling, thereby sensitizing resistant tumor cells. Consistent with this mechanism, previous work has shown that both normal and malignant granulosa cells secrete GDF15 into the culture medium, consistent with its role as a stress-responsive feedback signal^45^. While our findings point to such a mechanism, further studies will be needed to define the precise molecular events.

To examine the broader relevance of GDF15 in the tumor microenvironment, we used a recently published meta-analysis of lung adenocarcinoma tumor samples from The Cancer Genome Atlas^34^. Expression levels were classified as low or high using a median split, and their association with cytotoxic immune cell infiltration was compared among GDF15 and known necroptosis effectors (RIPK1, RIPK3, and MLKL). While higher expression of RIPK3, RIPK1, and MLKL correlated with increased CD8+ T cell infiltration, GDF15 displayed the opposite trend; elevated levels were associated with reduced infiltration (Fig. 4E). This observation aligns with our hypothesis that GDF15 acts as a negative regulator of necroptosis and may contribute to a reduced pro-inflammatory phenotype in the tumor microenvironment.

Taken together, our findings support a mechanistic model in which GDF15 acts as a RIPK3-interacting “brake”, suppressing downstream necroptotic signaling and limiting cell lysis. Loss or inhibition of GDF15 consistently results in increased necroptotic death, whereas its presence appears to restrain RIPK3-induced necroptotic signaling. By directly interacting with RIPK3, GDF15 appears to control the threshold for necroptotic activation in tumor cells, providing a potential link between cytokine signaling, inflammatory regulation, and programmed necrosis.

## DISCUSSION

Understanding and harnessing molecular signaling involved in programmed cell death has important therapeutic implications. Apoptosis is the most extensively studied form of programmed cell death, yet its therapeutic potential in cancer remains limited^46,47^. Many tumors acquire mechanisms to evade apoptosis, such as caspase inactivation or dysregulation of BCL-2 family proteins, which allow them to resist treatments that rely on apoptotic priming^47^. This widespread resistance underscores the need for alternative strategies. Necroptosis provides an alternative, caspase-independent route to cell death, triggered downstream of TNF receptor signaling. Upon TNF engagement, the assembly of distinct signaling complexes dictates whether cells activate NF-κB–driven survival, undergo caspase-dependent apoptosis, or commit to necroptosis when caspase activity is blocked^5,23^. This built-in plasticity of TNF signaling highlights necroptosis as a particularly attractive pathway to exploit, as it bypasses apoptotic resistance while simultaneously releasing inflammatory signals that can recruit and activate immune cells. Studying regulators of this pathway is therefore crucial to understand how tumor cells evade immunogenic cell death and to uncover opportunities for reprogramming TNF-induced cell death outcomes.

Our study identifies two previously unrecognized regulators of necroptotic cell death, TCOF1 and GDF15, using an unbiased RIPK3 interactome approach. By applying immunoprecipitation–mass spectrometry under active necroptotic conditions, we mapped RIPK3-specific protein–protein interactions and uncovered inhibitory nodes that had not been previously linked to necroptotic signaling. Functionally, loss of either TCOF1 or GDF15 markedly enhanced necroptotic cell death, positioning both proteins as negative interactors to this highly inflammatory form of programmed cell death.

While TCOF1 is primarily known for its role in ribosome biogenesis and craniofacial development, its inhibitory function in necroptosis suggests a previously unappreciated cytoplasmic activity in death signaling regulation^38^. Future work will be required to delineate whether TCOF1 influences necrosome assembly directly or modulates cellular stress pathways that converge on necroptosis. In contrast, GDF15 emerged not only as a functional suppressor but also as a direct binding partner of RIPK3. Our reciprocal immunoprecipitation experiments demonstrate that GDF15 directly interacts with RIPK3 and restrains necroptotic signaling. Our findings suggest that GDF15 is a cytokine that exerts dual control: acting intracellularly to restrain RIPK3 activity while also functioning extracellularly as part of a stress-responsive feedback loop. By binding RIPK3, GDF15 may function at the intersection of cell-intrinsic death regulation and extracellular immune modulation, thereby enabling tumors to suppress necroptotic signaling and subsequent immune activation.

The identification of GDF15 as a necroptosis suppressor opens opportunities for therapeutic intervention. Unlike RIPK1, RIPK3, and MLKL, whose elevated expression correlates with increased T cell immune infiltration, higher GDF15 levels were associated with reduced infiltration. This inverse relationship, consistent with our functional assays, suggests that GDF15 can restrain necroptotic signaling through direct interaction with RIPK3 and may contribute to an immunosuppressive tumor microenvironment. Neutralization of GDF15 sensitized tumor cells to necroptotic signaling in our system, highlighting the feasibility of targeting necroptosis brakes to amplify immunogenic cell death. This concept is further supported by recent work showing that GDF15 neutralization can overcome immune checkpoint therapy resistance in solid tumors, reinforcing its broader therapeutic relevance^34^.

Collectively, our work establishes TCOF1 and GDF15 as novel inhibitory regulators of necroptosis and positions GDF15 as a direct RIPK3-interacting checkpoint in cell death signaling. Beyond providing mechanistic insights, our study illustrates the power of proteomic interactome mapping to uncover actionable molecular targets. By revealing necroptosis-suppressive mechanisms, we lay the groundwork for approaches that may enhance tumor immunogenicity through selective rewiring of TNF-induced programmed cell death.

## LIMITATIONS OF THE STUDY

Our study relies on pharmacological treatment to induce necroptosis, which may not capture all physiological regulatory interactions. Furthermore, while our data firmly establish GDF15 as a RIPK3 interactor, the precise molecular interface and the context in which this interaction occurs require further elucidation. In vivo studies will also be critical to define the extent to which GDF15 regulates necroptosis within the tumor microenvironment, as this study is based on selected tumor lines. Lastly, phenotypic experiments employing Visugromab provide a pharmacological proof of concept, but additional validation is needed to firmly establish therapeutic relevance.

## ACKNOWLEDGMENTS

This work received support from the NWO-funded Netherlands Proteomics Center through the National Road Map for Large-scale Infrastructures program XOmics (Project 184.034.019).

## METHODS

### Cell culture

Human colorectal adenocarcinoma cells (HT-29), human breast adenocarcinoma cells (MDA-MB-231), human embryonic kidney cells (HEK293T), and human lung carcinoma epithelial cells (A549) were maintained in Dulbecco’s modified Eagle medium (DMEM; Gibco®; ThermoFisher Scientific, Waltham, MA, USA), supplemented with 10% fetal bovine serum (FBS; HyClone/GE, Malborough, MA, USA) and 1% penicillin/streptomycin (PenStrep, DE17-602E; Lonza, Basel, Switzerland). Regular mycoplasma testing was conducted to ensure contamination-free cultures. All cells were maintained in a humidified incubator at 37°C with 5% CO2. Experiments were conducted between the 5th and 10th passages to minimize cell heterogeneity.

### SgRNA construction and lentivirus production

For the construction of a sgRNA-expressing vector, DNA oligonucleotides were annealed and ligated into BsmBI-digested LentiCRISPRv2 plasmid. Target sgRNA oligonucleotide sequences are listed in Table S3. Functional knockouts were achieved by employing a puromycin-selectable variant of lentiCRISPR-v2 for each sgRNA.

Briefly, HEKT293T cells were used for lentivirus production containing mutation and SgRNA control constructs, or a GFP-positive control to determine transfection efficiency. Medium-containing pseudo-virus was harvested and concentrated 2- and 3-days post-transfection. Tumor cells were seeded in a 6-well plate in a seeding density of 2.5 × 10^4^ per well, where different virus volumes ranging from 2.5 to 15 μl were added. Transduction was enhanced using Polybrene® (Sigma-Aldrich, Merck KGaA, Darmstadt, Germany) to neutralize charge interactions and increase binding between the pseudoviral capsid and the cellular membrane. A lethal dose of 10 μg/ml of puromycin (Sigma-Aldrich) was added 3 days post-transduction. Medium with puromycin was refreshed every 2 days until puromycin-resistant colonies appeared.

### Drug treatment

To induce necroptosis, a combinatory drug treatment was applied to the cells (TBZ): z-VAD-FMK (InvivoGen, #tlrl-vad, San Diego, CA, USA) was applied at a working concentration of 0.2 µg/mL, along with Birinapant (MedChemExpress, #HY-16591, Monmouth Junction, NJ, USA) at 2 µM, followed by incubation for 1 hour at 37°C. Subsequently, TNF-α (PeproTech, #300-01A, Cranbury, NJ, USA) was added to a final concentration of 30 ng/mL. For the negative control condition (TBZN), Necrostatin-1 (InvivoGen, #inh-ncst1, San Diego, CA, USA) was added at a final working concentration of 40 µM. GDF15 neutralizing antibody Visugromab (MedChemExpress, #CTL-002) was added at a final working concentration of 10µg/ml.

### Cell cytotoxicity assay

For the cytotoxicity assay, HT-29 and A549 cells were seeded at 20,000 cells / well and MDA-MB231 cells at 30,000 cells / well in a 96-well plate and left to attach overnight in an incubator at 37°C, 5 % CO_2_. On the day of the experiment, the medium was replaced to media containing Incucyte® cytotox green dye (Sartorius, #4632, Göttingen, GER). The cells were then transferred to an Incucyte® SX1live-cell imaging system (Sartorius, Göttingen, GER) where they were scanned in 15-minute intervals for a period of 24hours post treatment.

### Immunoblotting

Cells were harvested and lysed in IP lysis buffer (30mM TRIS-HCl pH 7.4, 120mM NaCl, 2mM EDTA, 2mM KCl, 1% Triton X-100), containing 1x protease inhibitor (complete EDTAfree, #11873580001, Roche) and 1 x phosphatase inhibitor (PhosStop, #4906845001, Roche) on ice for 30 minutes. Samples were spun down in the centrifuge at 4°C for 10 minutes at max speed. The supernatant and pellet were separated, and the protein amount of the supernatant was determined using the Pierce BCA Protein Assay Kit (ThermoFisher Scientific, MA, USA). Cell lysates were denatured and reduced in XT Sample buffer (Bio-Rad) supplemented with 25 mM DTT at 95°C for 5 minutes. Proteins were then separated on a 12% Bis-Tris SDS-PAGE gel (Bio-Rad) and electroblotted onto PVDF membranes. The membranes were blocked with 5% non-fat milk (Nutricia, Zoetermeer, NL) in TBS 1X supplemented with 0.1% Tween 20 (TBS-T) and incubated overnight at 4°C with the following primary antibodies: anti-b-actin (1:1000, 4970S, Cell Signaling, Danvers, MA, USA), anti-GDF15 (1:1000, 79996S, Cell Signaling, Danvers, MA, USA), anti-MLKL (1:1000, 14993S, Cell Signaling, Danvers, MA, USA), anti-PMLKL (1:1000, 91689T, Cell Signaling, Danvers, MA, USA), anti-PRIPK1 (1:1000, 44590S, Cell Signaling, Danvers, MA, USA), anti-PRIPK3 (1:1000, 93654S, Cell Signaling, Danvers, MA, USA), anti-RIPK1 (1:1000, 3493T, Cell Signaling, Danvers, MA, USA), anti-RIPK3 (1:1000, 86671S, Cell Signaling, Danvers, MA, USA), anti-SEMG1 (1:1000, Abbexa, Cambridge, UK), anti-SEMG2 (0.1ug/mL, Abbexa, Cambridge, UK). As a secondary antibody, anti-rabbit IgG, HRP linked (1:1000, 7074S, Cell Signaling, Danvers, MA, USA) was used.

### Co-Immunoprecipitation and sample preparation

Cells were harvested and lysed in IP lysis buffer and centrifuged at 4°C for 10 minutes at max speed. The supernatant was then added to 25 µL of magnetic beads (IP kit, 88805, Pierce, ThermoFisher Scientific, Waltham, MA, USA), which were previously crosslinked to either anti-PRIPK3 (1:1000, 13526, Cell Signaling, Danvers, MA, USA), or anti-GDF15 (1:1000, 79996S, Cell Signaling, Danvers, MA, USA), according to the manufacturer’s protocol. The procedure was adapted and optimized from published methods to maximize recovery of RIPK3-associated complexes^48^. The mixture was incubated on a rotating platform at 4°C overnight. The following day, the beads were collected using a magnetic stand and washed five times with 500 µL of lysis/wash buffer (IP kit). To elute, 100 µL of elution buffer (0.5 M glycine, pH 2) was added to the beads and incubated for 5 minutes before being neutralized with 10 µL of neutralization buffer (Tris, pH 8.5). This step was repeated once more. A second elution with 5% SDS in 50 mM TEAB (pH 8.5) was conducted for maximum recovery and eluates were digested using S-TRAP microfilters (C02-micro-10, ProtiFi) as previously described^22,49^. Digested peptides were eluted and dried in a vacuum centrifuge before LC-MS analysis.

### LC-MS/MS

For mass spectrometry analysis, spectral data were acquired with an Orbitrap Exploris 480 mass spectrometer (Thermo Fisher Scientific, USA) coupled to Ultimate3000 (Thermo Fisher Scientific) UHPLC. Peptides were trapped on an Acclaim Pepmap 100 C18 (5 mm × 0.3 mm, 5 μm, ThermoFisher Scientific) trap column and separated on an in-house packed analytical column (75 µm, ReproSil-Pur C18-AQ 2.4 µm resin (Dr. Maisch GmbH). Solvent B consisted of 80% acetonitrile in 0.1% FA. Trapping of peptides was performed for 2 min in 9% B followed by peptide separation in the analytical column using a gradient of 13 to 44% B in 95 min. After peptide separation, gradients were followed by a steep increase to 99% B in 3 min, a 5-min wash in 99% B, and a 10-min re-equilibration at 9% B. Flow rate was kept at 300 nL/min. MS data was acquired using a DDA method at a resolution of 60,000 and a scan range of 375–1600 m/z. Automatic gain control (AGC) target was set to 3E6, under standard calculation of maximum injection time. The cycle time for MS2 fragmentation scans was set to 1 s. Peptides with charged states 2 to 10 were fragmented, with a dynamic exclusion of 16 s. Fragmentation was done using stepped higher-energy collisional dissociation (HCD)-normalized collision energy of 28%. Fragment ions were accumulated until a target value of 1 × 105 ions was reached or under a maximum injection time of 300 ms, with an isolation window of 1.4 m/z before injection in the Orbitrap for MS2 analysis at a resolution of 15,000.

### Data-dependent acquisition and statistical analysis

RAW files were processed with MaxQuant (version 2.6.3.0) and MS/MS spectra were searched against the UniProt human database (9606, Homo Sapiens) including common contaminants. Trypsin digestion and a maximum of three miscleavages were set using fixed carbamidomethyl modification, and the variable modifications oxidized methionine, protein N-terminal acetylation, and serine/threonine/tyrosine phosphorylation. A false discovery rate (FDR) of 1% was enforced across protein, peptide, and modification levels. Protein abundances were determined through label-free quantification with default settings. Database search outcomes underwent additional statistical analysis using Perseus (version 2.1.5.0).

Statistical comparisons included two-tailed Student’s T-tests for comparing two means, one-way ANOVA followed by Dunnett’s test for comparing multiple groups to a single control, and two-way ANOVA followed by Dunnett’s post-hoc test for comparing two factors. Specific exceptions to these methods are detailed in the corresponding figure legends. Statistical analyses and plots were conducted using Perseus (version 2.1.5.0), amica proteomics^50^, Prism 10 (Graphpad Software Inc., version 10.1.2), or in house R scripts. The statistical analysis of the phenotypic experimental data was performed using the Incucyte SX1 software, GraphPad Prism (Graphpad Software Inc., version 10.1.2), and in house R scripts. Results were shown as mean ± standard deviation (SD). Workflow figures were created partially with Biorender and bio-icons.com along with in-house illustrations.

---

**Supplementary Figure 1.**
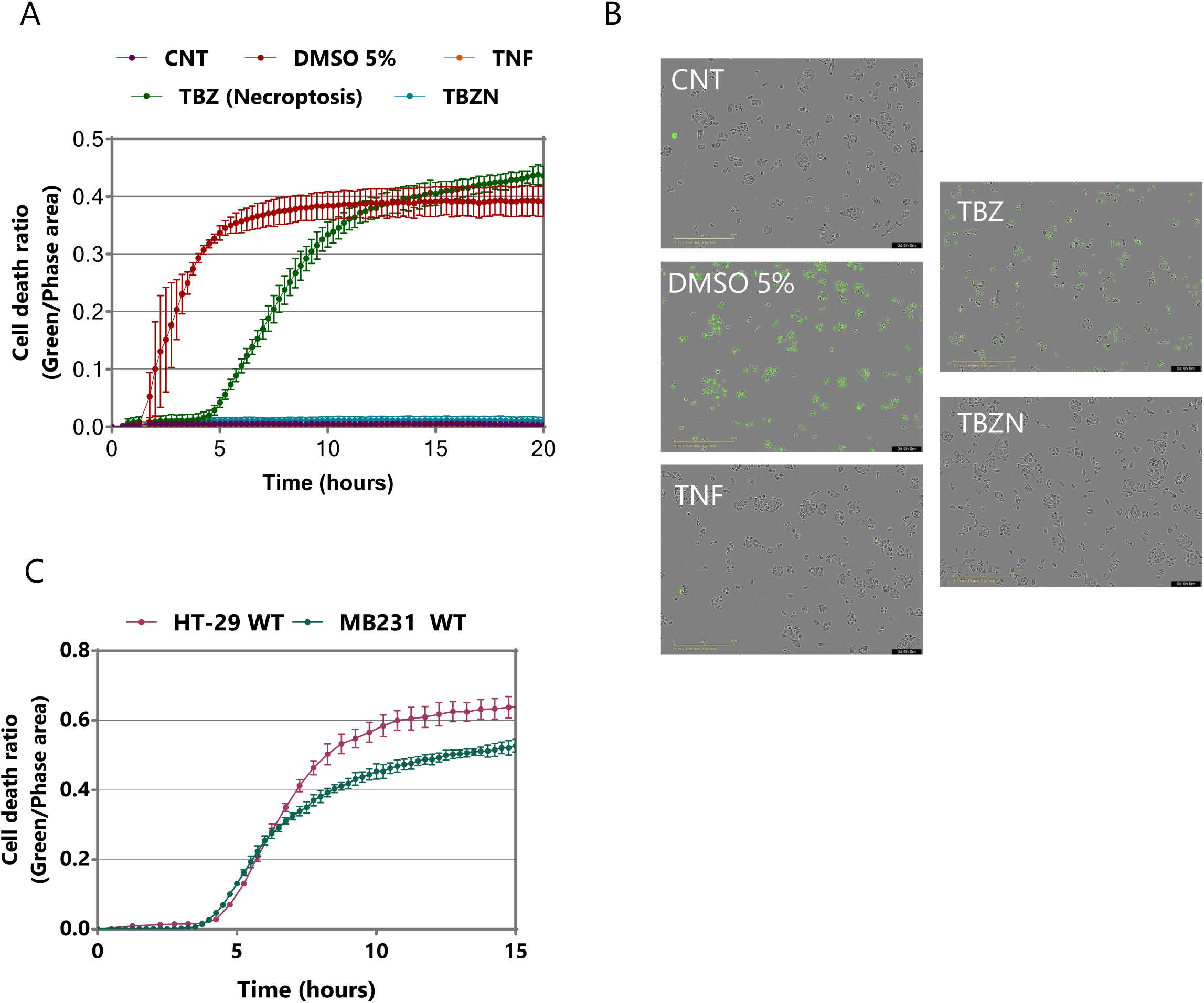
TNF-induced necroptosis is conserved across tumor types but shows lower sensitivity in MDA-MB231 cells. (A) Live-cell imaging of necroptotic death kinetics using MDA-MB231 cells under indicated treatments using Incucyte SX1 (Sartorius). Tumor cells were scanned in 15-minute intervals over 24 hours post treatment. Error bars represent standard deviation (SD) between replicates (n=3). Background signal was subtracted at the time of treatment (t=0) and green (cell death) signal is represented as a ratio to phase signal. **(B)** Representative images of MDA-MB231 cells following 6 hours of treatment. Scale bars: 400 µm. **(C)** Comparative sensitivity of HT-29 and MDA-MB231 cells to necroptotic treatment. Error bars represent standard deviation (SD) between replicates (n=3).

**Supplementary Figure 2.**
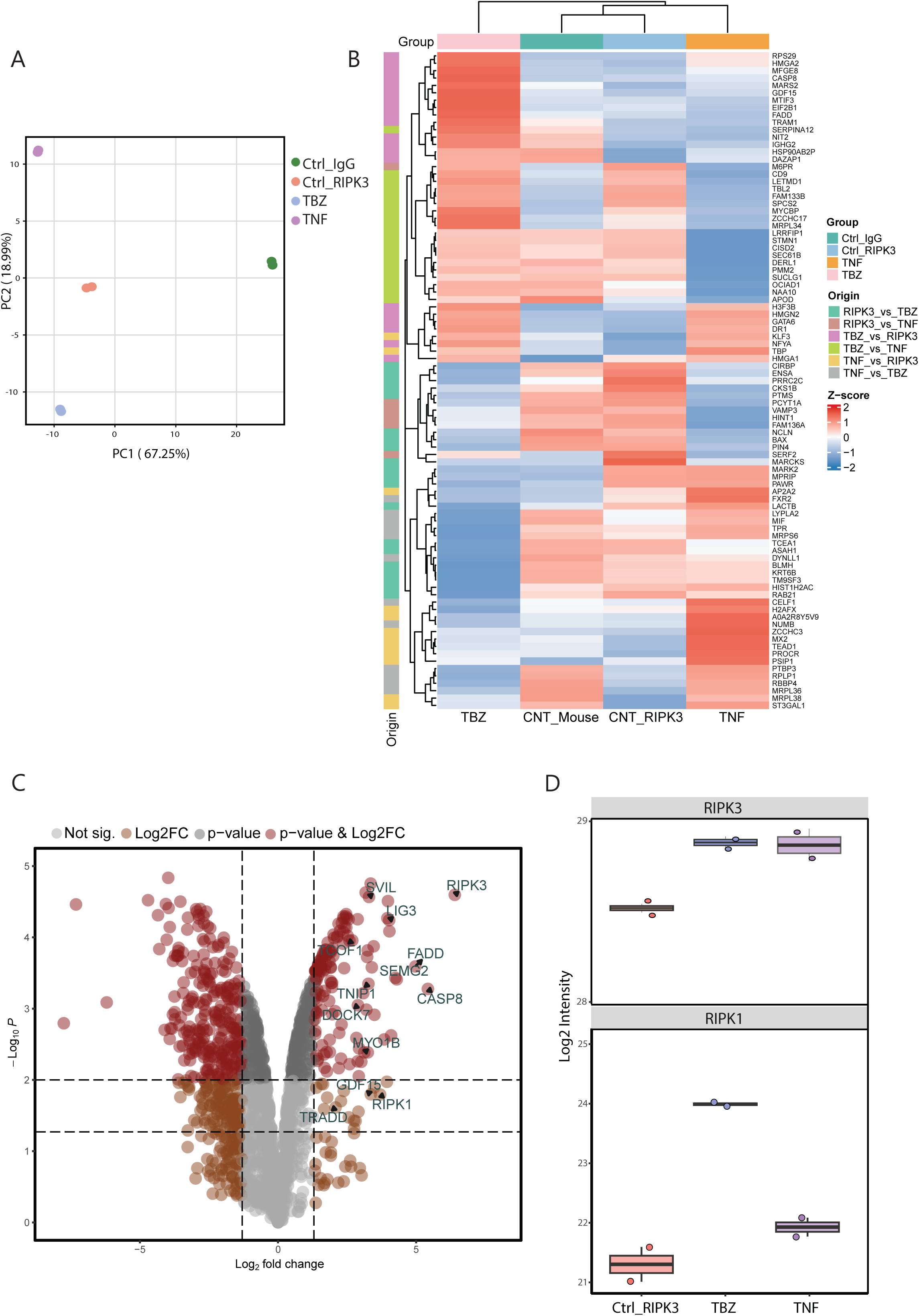
Identification of RIPK3 interactions through coIP-MS analysis. **(A)** Principal component analysis (PCA) of RIPK3 pulldowns under the indicated treatments, showing clear separation of TBZ-induced samples. **(B)** Relative abundance of the top significant hits from pairwise comparisons between treated and non-treated RIPK3 pull downs. Values are Log2 transformed, represent the average of 2 biological replicates per treatment and missing values were imputed. Color coding indicates treatment condition, and clustering reflects correlations between conditions. Z-score normalization was applied. **(C)** Relative abundance of TBZ-induced RIPK3 pull downs against IgG Ctrl in another independent experiment. Dashed lines indicate cutoffs at p < 0.05 and p < 0.01. **(D)** Relative abundance of RIPK3 and RIPK1 across treatment conditions. Values are Log2 transformed and represent the average of 2 biological replicates per treatment. IgG Ctrl is not shown as RIPK3 and RIPK1 were not detected in this condition.

Supplementary Table 3 Target sgRNA oligonucleotide sequences CRISPR-mediated knockouts.

**Supplemetary Figure 3.**
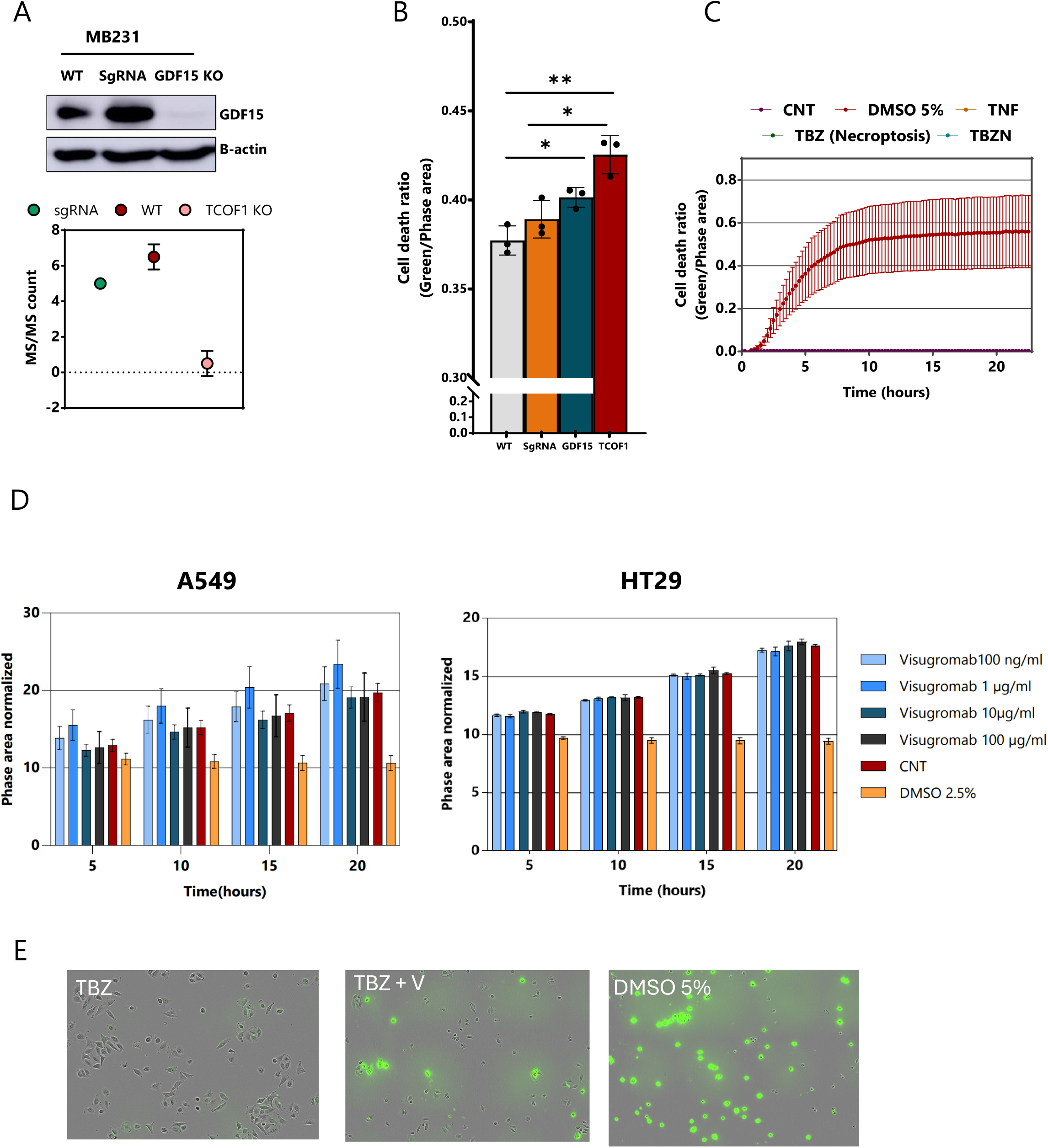
Functional roles of GDF15 and TCOF1 in MDA-MB231 cells and therapeutic targeting of GDF15 in A549 cells. **(B)** Validation of functional GDF15 knockout by Western blot (top) and quantification of MS/MS counts for WT, sgRNA control, and sgTCOF1 cells (bottom) in MDA-MB231 cells. **(B)** Quantification of necroptotic cell death in MDA-MB231 WT, sgRNA control, GDF15 KO, and TCOF1 KO cells following TBZ treatment. Measurements were taken at 20hrs post treatment. Data are mean ± SD (n=3); statistical significance was assessed by two-way ANOVA with Tukey’s multiple comparisons *p < 0.05, **p < 0.01, ***p < 0.001. **(C)** Live-cell imaging of necroptotic death kinetics under indicated treatments using Incucyte SX1 (Sartorius). A549 cells were scanned in 15-minute intervals over 24 hours post treatment. Error bars represent standard deviation (SD) between replicates (n=3). Background signal was subtracted at the time of treatment (t=0) and green (cell death) signal is represented as a ratio to phase signal. **(D)** Titration of Visugromab in A549 (left) and HT-29 (right) cells, showing no significant effects on cell viability in the absence of other treatments. Cells were treated with different Visugromab concentrations and cell growth was monitored with Incucyte SX1 over 20 hours. Data represent mean ± SD (n=3). A working concentration of 10ug/ml was selected. **(E)** Representative images corresponding to cell death rates of A549 cells following 9 hours of treatment with the GDF15 neutralizing antibody Visugromab in combination to necroptotic treatment. Scale bars: 400 µm.

